# SARS-CoV-2’s evolutionary capacity is mostly driven by host antiviral molecules

**DOI:** 10.1101/2023.04.07.536037

**Authors:** Kieran D. Lamb, Martha M. Luka, Megan Saathoff, Richard Orton, My Phan, Matthew Cotten, Ke Yuan, David L. Robertson

## Abstract

The COVID-19 pandemic has been characterised by sequential variant-specific waves shaped by viral, individual human and population factors. SARS-CoV-2 variants are defined by their unique combinations of mutations and there has been a clear adaptation to human infection since its emergence in 2019. Here we use machine learning models to identify shared signatures, i.e., common underlying mutational processes, and link these to the subset of mutations that define the variants of concern (VOCs). First, we examined the global SARS-CoV-2 genomes and associated metadata to determine how viral properties and public health measures have influenced the magnitude of waves, as measured by the number of infection cases, in different geographic locations using regression models. This analysis showed that, as expected, both public health measures and not virus properties alone are associated with the rise and fall of regional SARS-CoV-2 reported infection numbers. This impact varies geographically. We attribute this to intrinsic differences such as vaccine coverage, testing and sequencing capacity, and the effectiveness of government stringency. In terms of underlying evolutionary change, we used non-negative matrix factorisation to observe three distinct mutational signatures, unique in their substitution patterns and exposures from the SARS-CoV-2 genomes. Signatures 0, 1 and 3 were biased to C**→**T, T**→**C/A**→**G and G**→**T point mutations as would be expected of host antiviral molecules APOBEC, ADAR and ROS effects, respectively. We also observe a shift amidst the pandemic in relative mutational signature activity from predominantly APOBEC-like changes to an increasingly high proportion of changes consistent with ADAR editing. This could represent changes in how the virus and the host immune response interact, and indicates how SARS-CoV-2 may continue to accumulate mutations in the future. Linkage of the detected mutational signatures to the VOC defining amino acids substitutions indicates the majority of SARS-CoV-2’s evolutionary capacity is likely to be associated with the action of host antiviral molecules rather than virus replication errors.

## 1 Introduction

The COVID-19 pandemic began in late 2019 following a zoonotic spillover event of a SARS-related coronavirus, subsequently named SARS-CoV-2, in Wuhan city, China [1, 2]. The extensive and rapid global spread of this new human coronavirus and detrimental impact on human health has rendered it among the most significant pandemics in recent history [3]. Different geographical regions of the world have reported varied infection patterns that are attributed to differences in population demographics and health care systems, diverse government responses [4, 5], the emergence of more transmissible variants[6, 7] and other viral, human and population factors. Since its emergence, SARS-CoV-2 has undergone significant genetic change such that numerous variants, i.e., distinct genotypes, have been identified [8], many with altered phenotypic properties[9]. The World Health Organization (WHO) and other public health bodies have classified variants that pose an increased risk to global public health (due to increased transmissibility, increased virulence, or decrease in effectiveness of public health measures relative to 2019/early 2020 SARS-CoV-2 variants) as variants of concern (VOCs) and variants of interest (VOIs)[10]. The first SARS-CoV-2 variants to emerge in 2019 and then the more transmissible+S:D614G variant followed by the VOCs (Alpha, Beta, Gamma, Delta and currently Omicron) have driven significant and sequential waves of SARS-CoV-2 infections internationally.

Contrary to geographical variation in emergence of SARS-CoV-2 variants [11–14], we also witness commonality of mutations across independent variants[15], indicating convergent evolution[16]. Viral mutations arise from a diverse set of processes (viral polymerase replication errors mistakes, host anti-viral editing processes, etc.) which can be identified by the characteristic mutational signatures that they leave on the genome [17, 18]. Such characterisation of dominant mutational processes is routinely used in cancer genomics [19]. The catalogue of SARS-CoV-2 nucleotide changes show clear mutational patterns suggestive of a role for host mutational processes in introducing changes in the viral RNA [20, 21]. These potentially dominate in SARS-CoV-2 evolution due to the action of a proofreading enzyme such that point mutations introduced in replication are corrected.

The generation of virus diversity, the key to virus persistence by generating novel variants and thus evolutionary capacity, is multi-faceted [22], yet our understanding of the relative importance of underlying mutational processes linked to the action of host anti-viral molecules is still very limited. Given that SARS-CoV-2 continues to develop novel variants, many associated with sets of previously observed (convergent) or new beneficial mutations, it is critical that we improve our understanding of the mechanisms and source of evolutionary change.

Along with routine surveillance of SARS-CoV-2 infections, there has been an unprecedented global sequencing effort resulting in databases containing many millions of genome sequences, in particular GISAID [23]. Here we examined this data to describe the global molecular epidemiology and evolution of SARS-CoV-2. Using regression models we first examined how viral properties and public health measures have influenced the magnitude of infection waves in different geographic locations. Satisfied that SARS-CoV-2 variants have been an important driver of infections we then used non-negative matrix factorisation to characterise the mutational processes involved in the generation of variants and their changing patterns of activity over time.

## 2 Results

### 2.1 Characterising the SARS-CoV-2 Waves Regionally

This first part of the study reports on global SARS-CoV-2 data from 24/12/2019 to 28/01/2022 only as limited public health measures were in place after this time. We observed 1,544 distinct SARS-CoV-2 lineages from 7,348,178 sequences. 88% of the infections in the global pandemic during this time frame) were caused by a subset of 13 Pango lineages (**Supplementary Table A2**). There was varied geographical diversification of the virus, and different lineages displaced one another over time.

A clear dominance of a subset of variants and replacement of these through time was observed. This “wave” infection pattern was evident in all geographic locations. Although biased by testing rates, Europe and the Americas had the highest infection rates, reporting up to 450 cases per million population per day (**Figure 1**). The emergence or introduction of VOCs coincided with a steep increase in infection rates globally. For example, cases in Asia showed a steep rise in February 2021, which peaked in May 2021 (**Figure 1 panel Asia**). During this period, Alpha and Delta comprised greater than 75% of the SARS-CoV-2 virus reported in the sequence data. Africa and Oceania on the other hand displayed overall sustained low case numbers. Despite this, Beta dominated the second wave in parts of Africa while Alpha dominated the third Oceanic wave. After its emergence in March 2021, Delta spread to become the predominant lineage across all the continents, **Figure 1**.

**Fig. 1.**
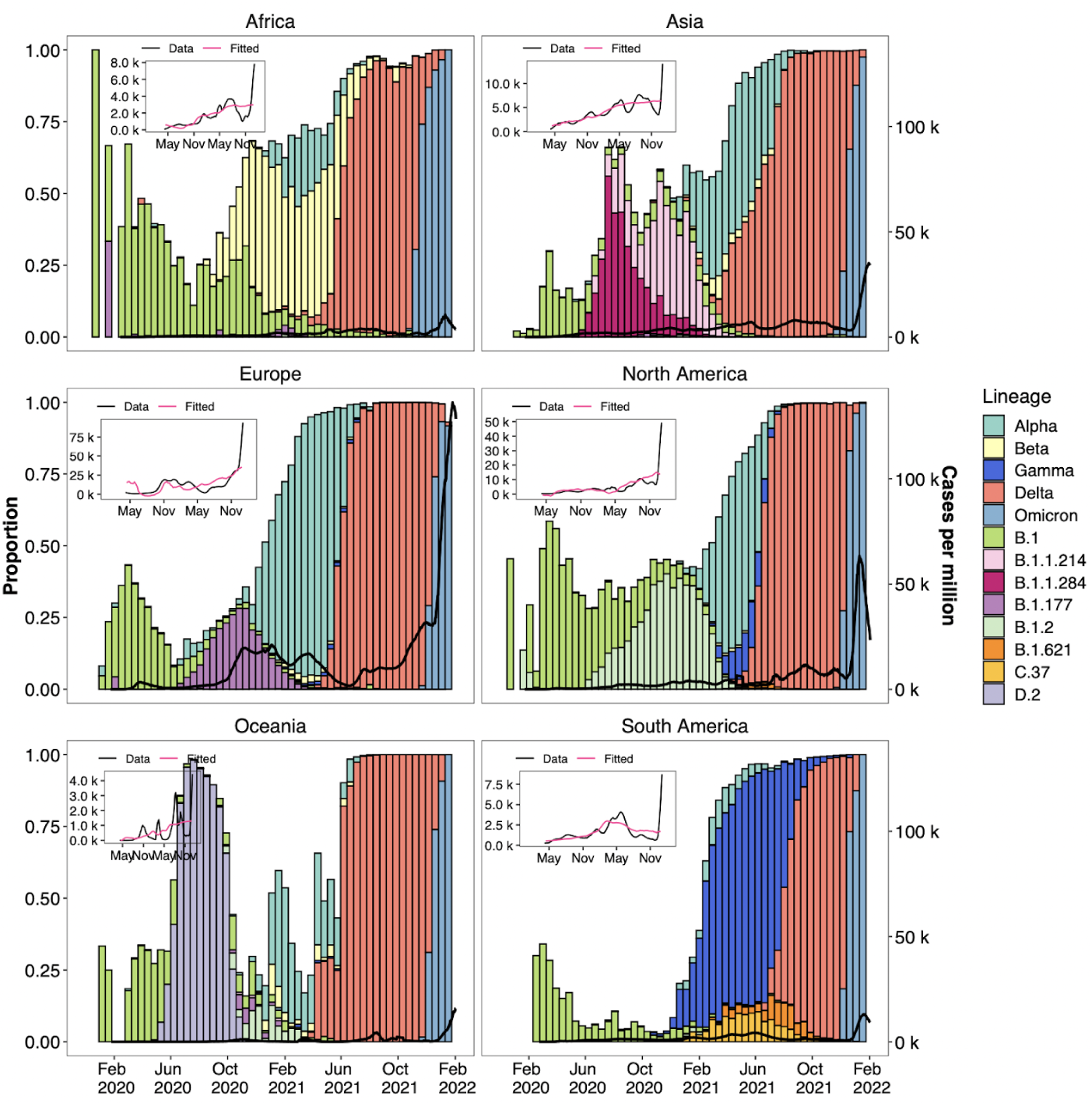
Continent-level SARS-CoV-2 lineage dynamics and pandemic curves. **(Main)** Dark solid lines show a 14-day rolling average of reported SARS-CoV-2 cases. Solid bars show the biweekly proportions of common lineages. Bars are coloured by lineage. **(Inset)** Model fitting through multiple linear regression. Black solid lines show a 14-day rolling average of reported SARS-CoV-2 cases. Pink solid lines show fitted mean response values of infection rates with predictor values as input.

### 2.2 Covariates of the waves

We investigated the degree to which public health measures and viral properties explain continent-specific reported cases of infection. Correlation analysis at the global level showed a significant correlation between infection rates and the four predictor variables (government stringency, vaccination, virus diversity and fitness),**Supplementary Table A3**.

Regression analysis shows that the impact of predictor variables on the magnitude of reported cases were found across all continents, p-value less than 1.804e-07. We defined no significance as p-value greater than 0.05, weak significance as p-value between 0.05 and 0.001, and high significance as p-value less than 0.001. Our analysis showed that, in the presence of the other variables, government stringency had no significant impact in South America, a weakly significant impact in Europe and a strongly significant impact on reported cases in Africa, Asia, Oceania and North America. Second, vaccination had a strongly significant impact in all continents except Asia and Europe, where the significance was low. Third, virus fitness was associated with high infection numbers in all continents except Africa (no significance). Fourth, virus diversity was associated with high infection numbers in Oceania, North and South America, and showed no association in Africa, Asia and Europe. Overall, R squared varied from 0.17 in Europe to 0.87 in South America. The predictor variables best explained the cases of infection in Africa, Asia, Oceania and South America (R-squared greater than 0.5), and less so in Europe and North America (R-squared less than 0.5), **Supplementary Table A1**. The fitted mean response values with predictor values as the input resembled the rise and fall of infection cases, **Figure 1**.

For country-level analysis, we included 17 countries from six continents based on the completeness of data (availability of sequence data in every 14 day bin). Pandemic plots were visualised using biweekly bins and multiple linear regression was fitted using the same approach. Different countries had varying lineage dynamics as illustrated in **Supplementary Figure A1**. The four predictor variables had varying impacts on infection rates across countries, **Supplementary Figure A2**. Despite some differences related to the population level processes investigated here, there is a clear variant replacement process taking place. As the generation of novel variants is fundamentally a mutation dependent process we next investigated the underlying patterns of mutations being generated through time.

### 2.3 Identifying putative mutational processes contributing to changes in SARS-CoV-2

New variants of concern have displaced viral lineages that were previously dominant in the population in different geographical regions and in some cases globally **Figure 1**. This behaviour has been observed with the original variants of concern (Alpha, Beta, Gamma) and then globally with the Delta and Omicron lineages. We decided to investigate whether these variant “wave” events (periods of time where infections are driven by a single variant or variant family i.e VOC) were driven by the activity of mutational processes. Each of the variants are mutationally distinct from the other, having acquired mutations that make them independent. Detecting the patterns of mutations in the data allows us to observe which processes are most active and could be driving the emergence of variants.

Mutations were called using constructed references for each of the Pangolin lineages which we call tree-based referencing. The full alignment of 13,278,844 sequences up to 26/10/2022 was used. Of those 13 million sequences 2,195,182 sequences were selected as they contained 5,726,144 newly arisen mutations. Cytosine to thymine mutations were the most common and were the primary substitution category for most weeks where sequences were recorded.

3 signatures were identified with distinct substitution patterns using Non-Negative Matrix Factorisation (NMF). Signature 1 is heavily biased towards cytosine to thymine (C→T) mutations, which is a substitution class often related to cytidine deamination via the activity of APOBEC enzymes [20, 24]. Note, SARS-CoV-2 has an RNA genome but we refer to Uracil as a Thymine to match pre-existing DNA mutational signature notations. The Signature had a high probability of ACA, ACT and TCT contexts (adjacent nucleotides in the 5’ and 3’ direction of the mutated site), consistent with what was earlier reported by [20] as highly mutated contexts for C→T substitutions in SARS-CoV-2. Signature 2 is predominantly adenine to guanine (A→G), guanine to adenine (G→A) and thymine to cytosine mutations (T→C). The proportion of A→G and T→C mutations is approximately equal in this signature, which is indicative of a double-stranded mutational process like adenosine deamination via the activity of ADAR. SARS-CoV-2 mutations at adenine positions on the negative strand will be counted as thymine mutations due to the negative strand being used to replicate positive sense RNA, with the mutated A→G now pairing with a cytosine on the +sense RNA and replacing the original thymine [25, 26]. Signature 3 is predominantly composed of guanine to thymine (G→T) substitutions, a pattern that is thought to be induced by the activity of Reactive Oxygen Species(ROS) causing oxidation of vulnerable guanine bases [27, 28].

Signatures 1, 2 and 3 are biased to C→T, A→G/T→C and G→T substitutions as would be expected of APOBEC, ADAR and ROS respectively. Analysis from [29] also detected similar signature patterns from intra-host SARS-COV-2 sequence samples.

### 2.4 The dynamics of mutational processes through the pandemic

By using the available SARS-CoV-2 sequences we can measure the mutational signature activity across time as long as our samples are aggregated using time series annotations. Signature exposures (**Figures 4**) show that the APOBEC-like signatures remained the most prominent signature throughout the pandemic, although following the emergence of ADAR-like signature its activity reduced proportionally. The ADAR-like signature appears inconsistently in the early weeks, before establishing itself as a dominant signature after December 2020. It continues to expand after October 2021, just prior to the emergence of the Delta VOC. The ROS-like signature is by far the least active of the 3 but remains consistent until after January-February 2022 when it begins to drop to almost zero. This is around the time Omicron begins to emerge as the dominant VOC.

Combined signature activity reached a peak at week 86(**Figure 4 (b)**) while the number of unique mutations peaks shortly before at week 84(**Figure 5 (a) and (b)**). This is around the time the mutational signature dynamics appear to be shifting, with the ADAR-like signature contributing more unique mutations. We can see that this also coincides with the Delta VOC wave, which between May 2021 and January 2022 was the lineage group showing the greatest number of newly acquired mutations (**Figure 5**). Delta was the first VOC to dominate on a worldwide scale, out-competing other VOC’s like Alpha, Beta, and Gamma in their regions of origin. Omicron similarly repeated this phenomenon, almost entirely wiping out Delta globally within weeks of its emergence (**Figure 5 (b)**). We also see a marked decrease in the activity of the ROS-like signature following Omicron’s establishment as the dominant variant. This marks a change from Delta, which saw an increase in the ROS-like signature following its emergence. This becomes particularly apparent when we begin to look at signature activities within lineage subsets of the data.

#### 2.4.1 Signatures Dynamics Spatially and Variant-wise

After observing changes in signature activity during transitions between variants, we next investigated the differences between signature activities in variant-only subsets of the data as well as in continent subsets. We used the globally extracted signatures to extract exposures from the subsets using a Non-Negative Least Squares regression to retain the non-negativity constraint. This allowed for the measurement of signature activity in each of the subsets of interest.

The APOBEC-like signature was the most active in almost all the subsets as was expected from the global activity. The ROS-like signature was most active in the Delta subset as well as during the Delta wave in the continent subsets(**Figure 5**). The Non-VOC, Beta, and Omicron subsets appear to be least impacted by the ROS-like signature with almost zero activity in Omicron. The ADAR-like signature also shows low activity in the Non-VOC subset but is very active in the other VOC subsets in particular Alpha where it appears to be the most active process, overtaking the APOBEC-like process.

Continent subsets also consistently showed high activity of the APOBEC-like signature. The ADAR-like signature begins to consistently appear in all continents after 2020, with only small bursts of activity being detected before (**Figure 5 (d)**). This is again consistent with what we see in the global data. The ROS-like activity also mimics the global activity, appearing most prominently during the Delta wave.

### 2.5 Bridging the gap between signatures and amino acid mutations

Stratifying non-synonymous changes by their putative processes should provide insights into how mutational processes affect viral proteins. The 3 mutational processes that were recovered can be categorised by their dominant substitution types: APOBEC-like substitutions include CT and GA, ROS-like changes are represented by GT and CA changes, and ADAR-like changes are AG and TC substitutions. The majority of non-synonymous changes appear to be derived from mutational processes based on this substitution type matching (**Figure 6**). APOBEC-like changes are still the most frequent, with ADAR mutations coming in second and ROS mutations in third. Stratifying non-synonymous changes by their putative processes might provide insights into how mutational processes affect viral proteins. The 3 mutational processes that were recovered can be categorised by their dominant substitution types: APOBEC-like substitutions include CT and GA, ROS-like changes are represented by GT and CA changes, and ADAR-like changes are AG and TC substitutions. The majority of non-synonymous changes appear to be derived from mutational processes based on this substitution type matching. APOBEC changes are still the most frequent, with ADAR mutations coming in second and ROS mutations in third.

Using the tree-based references, we can also look at individual lineage reference sequences to observe which mutational processes have produced their mutations. The tree references were used since they are equivalent to a high-quality representative sequence and because many of the early real sequences contain sequencing errors.

The Alpha VOC tree-based reference contains 13 APOBEC-like changes, 4 ROS-like changes, and 4 ADAR-like changes. APOBEC-like changes accounts for 46% of all substitution mutations within the Alpha tree lineage sequence, with 53% of these mutations being non-synonymous changes. Clearly, this APOBEC-like process was frequently active prior to the Alpha VOCs emergence. The activity plots (**Figure 4**) show that this was the case for much of the pandemic, particularly prior to the Alpha lineage emergence around September 2020. It should be noted that while APOBEC-like mutations are by far the most frequent, only one is found within the Spike protein (producing the S:T716I change). ROS-like changes on the other hand were all non-synonymous, with 2 appearing in the Spike glycoprotein, including S:P681H and S:A570D. ADAR-like mutations were non-synonymous 75% of the time, with the only Spike mutation relating to the process being S:D614G which is present within all known variants of concern.

The Beta lineage emerged around the same time as Alpha (Autumn 2020) but has a smaller set of mutations. A greater proportion of the APOBEC mutations are non-synonymous in Beta (70%) including S:E484K which is reported to help the virus evade neutralising antibodies [30]. ADAR-like mutations resulted in S:D614G and S:D215G in the Spike coding region with one additional ADAR mutation in ORF1a/b. ROS-like mutations produced S:K417N in spike which is also reported to aid in antibody evasion [30, 31] like S:E484K. Gamma also emerged in Autumn 2020 and has 33 different mutations.

APOBEC-like mutations account for 15 of these with 2/3 being non-synonymous. Most of these changes are located in ORF1a/b, however, 5 are present in Spike including S:L18F, S:P26S, S:E484K, S:H655Y and S:T1027I. ADAR-like changes resulted in fewer amino acid changes in Gamma than in other VOCs, with only 40% of changes being non-synonymous (one of which is S:D614G). ROS mutations in the Gamma lineage were all non-synonymous, with 4 of the 5 mutations in Spike.

Delta was the first VOC to dominate worldwide and replace almost every other lineage in all regions. The original Delta sequence (B.1.617.2) contains 9 APOBEC-like mutations. 77% of these changes were non-synonymous and only 2 occurring within Spike. ADAR-like mutations were all non-synonymous and displaced throughout the virus ORFs including S, M, ORF7a and N. ROS changes in Delta are found in Non-Coding regions as well as the N and S ORFs. The only ROS-like change in Spike causes the S:T478K mutation.

Omicron is the most recent VOC to emerge, quickly replacing Delta globally. Omicron differs from earlier VOCs with a much greater density of Spike mutations relative to the other ORFs. The first identified Omicron lineage B.1.1.529 has 40 substitution mutations of which 32 are non-synonymous substitution changes. This is almost double that of Delta which only had 18. 5 of these substitutions were APOBEC-like changes, 5 were ROS-like changes and 5 were ADAR-like changes. Only 6 could not be attributed to one of the putative mutational processes. There are 4 non-synonymous ORF1a/b mutations despite this ORF being substantially larger than SARS-CoV-2’s other ORFs. Only one Spike substitution was synonymous out of the 21 total changes.

This number is even greater when looking at the major Omicron variants BA.1 and BA.2. BA.1 had 31 non-synonymous changes in Spike alone while BA.2 had 28. Between these 3 Omicron variants, only 2 Spike substitutions are non-synonymous out of a total of 40. 14 of the 40 changes are APOBEC-like, 8 are ROS-like and 7 are ADAR-like. This means approximately 29/40 of the changes appear to come from these three mutational processes. 20 of the 40 mutations observed in these variants were present in the RBD of Omicron, with 9 of these mutations thought to help Omicron evade the immune response or increase its transmissibility [32]. Of these beneficial RBD changes, 3 are potentially the result of APOBEC activity, 5 are ADAR and none are ROS. The high density of ADAR RBD mutations in a variant that has emerged as the ADAR signature has increased may suggest that the ADAR mutational process has driven the emergence of the Omicron variant.

#### 2.5.1 Signature Exposures and Highly Mutated Sequences in Wastewater Data

Similar trends, over time, in exposures are seen when ADAR-like, APOBEC-like, and ROS-like signatures are applied to publicly available wastewater data. Although the trend is seen at a lower resolution than global data, the ROS-like and APOBEC-like signatures dissipate over time in place of the ADAR-like signature (**Figure 7**). Although, the ADAR-like signature is not quite as strong as in the global data (**Figure 4, 7**). This suggests trends in mutational processes can be monitored using wastewater, not only mass sequencing of the population. Additionally, at time periods where a high level of virus diversity is expected, there are highly mutated sequences present in the wastewater (**Figure 7**). This suggest cryptic sequences in wastewater may be used to observe potential upcoming variants, similar to how known sequences have been back-traced to particular buildings using wastewater [33].

As chronic infections are implicated as a major contributor to VOC evolution [34, 35], it may be possible to parse highly-mutated cryptic sequences of interest from chronic infections out of wastewater data in the interest of detecting potential VOCs. Unfortunately, this is problematic to deconvolute as robust sequencing data for exclusively immunocompromised or chronically infected individuals is sparse. When sequences from known immunocompromised individuals are examined, the distribution of mutation types is consistent with global data, with APOBEC-like mutations being the most common, as expected for samples from January 2022 (**Figure 7**). Although this is less than conclusive, it underpins interesting avenues of investigation as to where, and with whom SARS-CoV-2 is evolving.

## 3 Discussion

In this study, we described SARS-CoV-2 lineage dynamics and identified temporal variables that are associated with increased numbers of infection cases. Both public health measures and virus properties were associated with the rise and fall of regional SARS-CoV-2 infection cases. These predictors have a varying impact across geographical locations. The continual mutation and emergence of new lineages are expected. As more of the global population’s immune system becomes sensitised to SARS-CoV-2, either through previous infection or vaccination, the virus undergoes viral escape in search of new fitness landscapes to ensure survival. In some regions, government stringency had limited significant impact on patterns of infection. This could be due to differences in implementation strategies and support, other competing predictor variables, as well as behavioural changes in citizens as a response to the restrictions. Our analysis showed that vaccination had a significant impact on the pattern of reported cases across all continents. This is despite the low vaccination coverage in some regions such as the African continent. Neither virus fitness nor diversity had a significant impact on reported cases in Africa, while virus diversity showed no significant impact in Asia. Africa has experienced low-level sustained transmission, speculated to be associated with low median population age, low population density, immune priming by the high prevalence of other infectious diseases and low testing capacity[36]. The absence of impact from viral diversity on reported cases in Asia can be explained by the emergence of the Delta variant. Delta, first identified in Asia, displaced other lineages to become the predominant lineage. Overall, the predictor variables best explained the reported cases in Africa, Asia, Oceania and South America, and less so in Europe and North America. We interpret this to be due to intrinsic differences such as vaccine coverage, testing and sequencing capacity, and the effectiveness of government stringency.

The extracted signatures from the global SARS-CoV-2 dataset show clear patterns describing mutational processes acting on the viral genome. The most prominent of these signatures Signature 1 (**Figure 3**) shows a marked bias towards C→T mutations, a signal indicative of the APOBEC family of cytidine deaminases [20, 21]. APOBEC enzymes have been shown to cause extensive C→T editing of DNA and RNA in human and viral genomes. However it is not yet clear whether they are the cause of this pronounced C→T bias in SARS-CoV-2 despite a number of other studies also observing other APOBEC-like mutational patterns [29, 37, 38]. Cytosines flanked by either an adenine or thymine in both the 3’ and 5’ direction appear to be the most pronounced targets of Signature 1. APOBEC editing was shown to have contexts outside of the traditional TpC when structural features of the nucleic acid such as hairpin loops are present [39]. Outside of structural features, APOBEC3A is thought to be the predominant cause for TpC changes, and is found to be expressed in lung tissue [40]. ApC changes are thought to be caused by APOBEC1, which in cell models was shown to efficiently edit the SARS-CoV-2 viral RNA [40]. APOBEC1 is found predominately in the Liver and small intestine, which are tissues also thought to be infected by SARS-CoV-2 [40, 41].

**Fig. 2.**
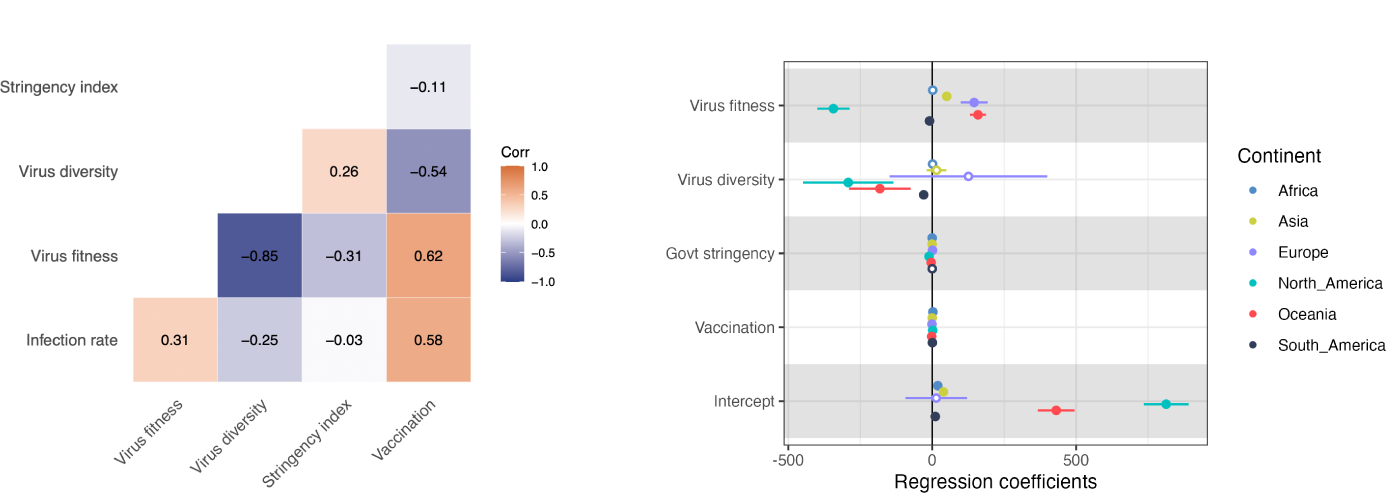
(a) Pearson’s correlation matrix of infection rate and predictor variables globally. Positive correlations are denoted in orange and negative correlations in blue, and colour intensity is directly proportional to coefficient value. (b) Forest plot of regression coefficients (95% confidence interval) for the association of infection rates and public health measures and viral properties. Circles are grouped by variables and coloured by continent. Hollow points indicate non-significant coefficients (p-value *>*0.05).

**Fig. 3.**
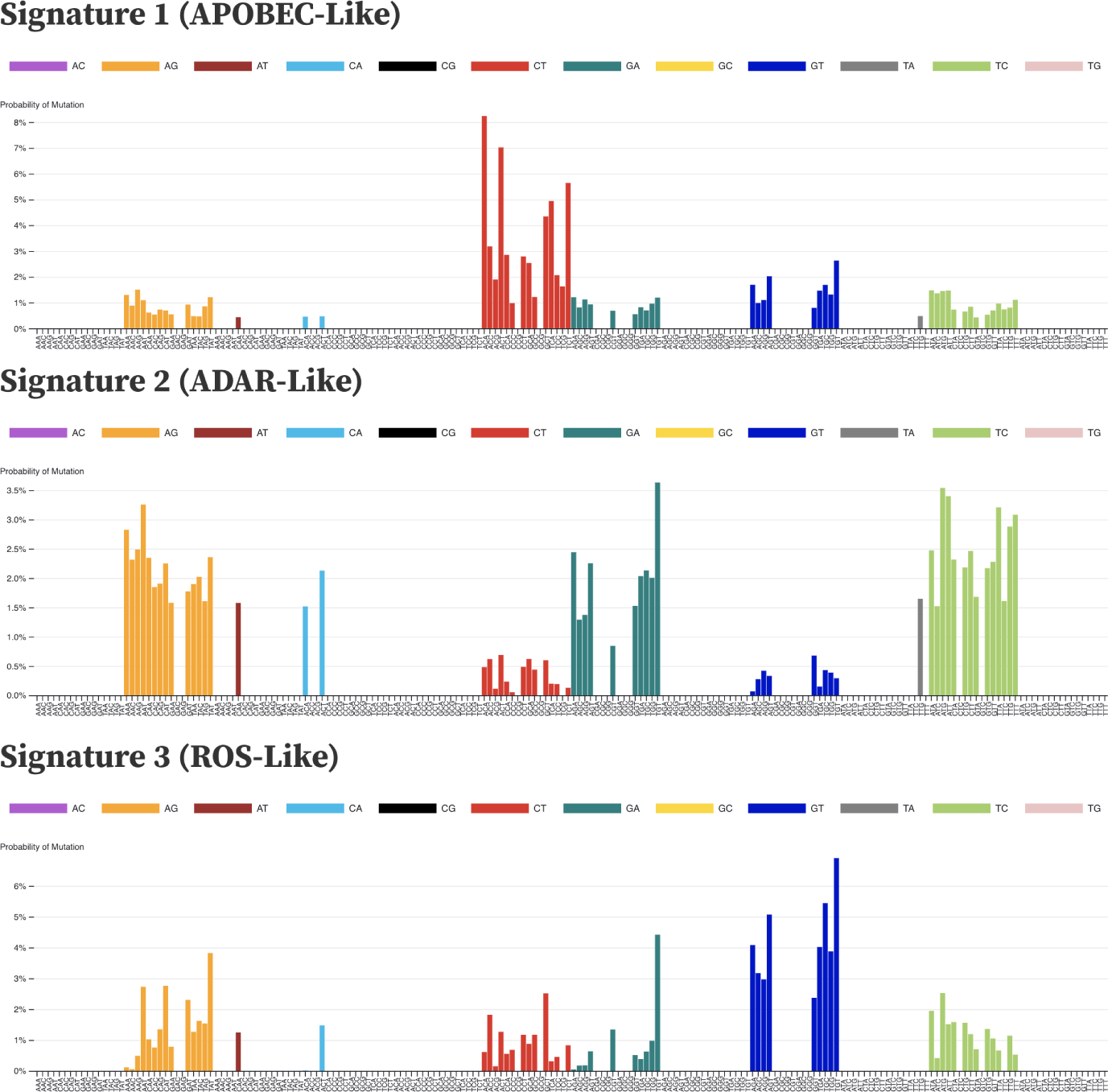
Mutational signatures extracted from the SARS-CoV-2 viral genomes by non-negative matrix factorisation. Signatures are patterns of probabilities for each category of substitution in a 3 nucleotide context. Each bar represents a context and is coloured by the substitution category of the mutation that occurs there. Each signature may represent a distinct mutational process. Signature 1 from SARS-CoV-2. Signature 1 is heavily biased towards cytosine to thymine (C→T) mutations, particularly in ACA, ACT and TCT contexts (consistent with what was earlier reported by [20]). Signature 2 from SARS-CoV-2 is predominantly adenine to guanine (A→G), Guanine to adenine (G→A) and thymine to cytosine mutations (T→C). Signature 3 is strongly guanine to thymine (G→T), a pattern that is thought to be caused by action of guanine oxidation by reactive oxygen species.

Signature 2 (**Figure 3**) has a nearly identical proportion of A→G and T→C mutations which are a known target of the ADAR family of adenine deaminases. ADAR enzymes typically operate on double-stranded RNA and convert adenine into inosine [25, 26]. Inosine forms base pairs with cytosine, which after another round of replication causes guanine to replace the inosine and complete the A→G change. As ADAR operates on both strands of dsRNA, the mutational signature resulting from the process is expected to contain an equal proportion of A→G and T→C mutations which is the case for Signature 2 [25]. Signature 2 also contains a number of G→A mutations, which may be caused by low-level APOBEC activity on the negative sense RNA strand. Since APOBEC only operates on the ssRNA, it is less likely to cause G→A substitutions than C→T due to the cellular strand bias present between the + and - sense RNA [38]. The negative strand will only be present during the replication phase of the virus while the positive strand will be present both on cell entry and on exit as the new viral particles are packaged to infect further cells. This is possibly why the negative sense APOBEC signature is present in Signature 2 since it may be operating at a similar level to ADAR on the negative strand.

Signature 3 (**Figure 3**) is dominated by G→T substitutions which are thought to be caused by reactive oxygen species (ROS) in the cell. Increases in oxidative stress as part of a ROS ‘burst’ have been associated with viruses during the early stages of infection [28, 29]. Guanine nucleotides are known to be vulnerable to oxidation, with the product 7,8-dihydro-8-oxo-2’-deoxyguanine (oxoguanine) pairing with adenine bases rather than cytosine [27, 28]. Similar to inosine causing A→G changes, this change to oxoguanine will result in a G→T mutation after a replication cycle. The lack of C→A changes in the signature also suggests that the mechanism is most active on the +ssRNA rather than the -ssRNA. The initial +ssRNA is found in the cytoplasm, meaning it can be easily accessed by ROS and other mechanisms of mutation. Viral replication is thought to take place within membrane-bound environments that aim to protect the RNA. The presence of dsRNA within these environments strongly suggests that this is the case [42] and may explain the relative lack of negative strand mutations in SARS-CoV-2 signatures.

Signature activities clearly change in both the global dataset and in the various subsets of the data for VOCs and Continents. In the global data (**Figure 4**) the APOBEC-like signature is dominant throughout the pandemic. The ADAR-like signature only begins to appear around November 2020, after which it appears consistently active for the remainder of the pandemic. This is approximately when variant of concern lineages began to emerge, as well as the beginning of the first vaccine rollouts. This is particularly apparent in the Alpha data subset where the ADAR-like signature is the most highly active mutational process (**Figure 5**). This pattern is not observed in the other VOC datasets, although Delta and Omicron have a large level of ADAR-like exposure as well despite the APOBEC-like signature remaining the dominant process in those subsets. The ROS-like signature appears to be most prominently found in the Delta lineage subset, and remains consistently at low levels in the global data until February 2022 when it appears to disappear almost entirely. The Omicron subset has little to no exposure of the ROS-like signature, and this happens to be the VOC almost exclusively circulating after February 2022. Why Omicron appears to have so little ROS-like exposure is unclear, although unlike previous VOCs Omicron differs in its preference of cell entry mechanism. Previous variants of the virus typically enter the cell using membrane fusion, where the viral membrane fuses with the cell membrane via the action of ACE-2 receptor binding and TMPRSS2 cleavage of the spike protein. Omicron instead favours an endosomal route of entry whereby the viral particle binds to the cell using ACE-2 and is enveloped by endocytosis into the cell. Cleavage of the spike protein then occurs via the action of Cathepsin L. which allows for the release of the viral RNA into the cytoplasm of the now-infected cell [43, 44].

**Fig. 4.**
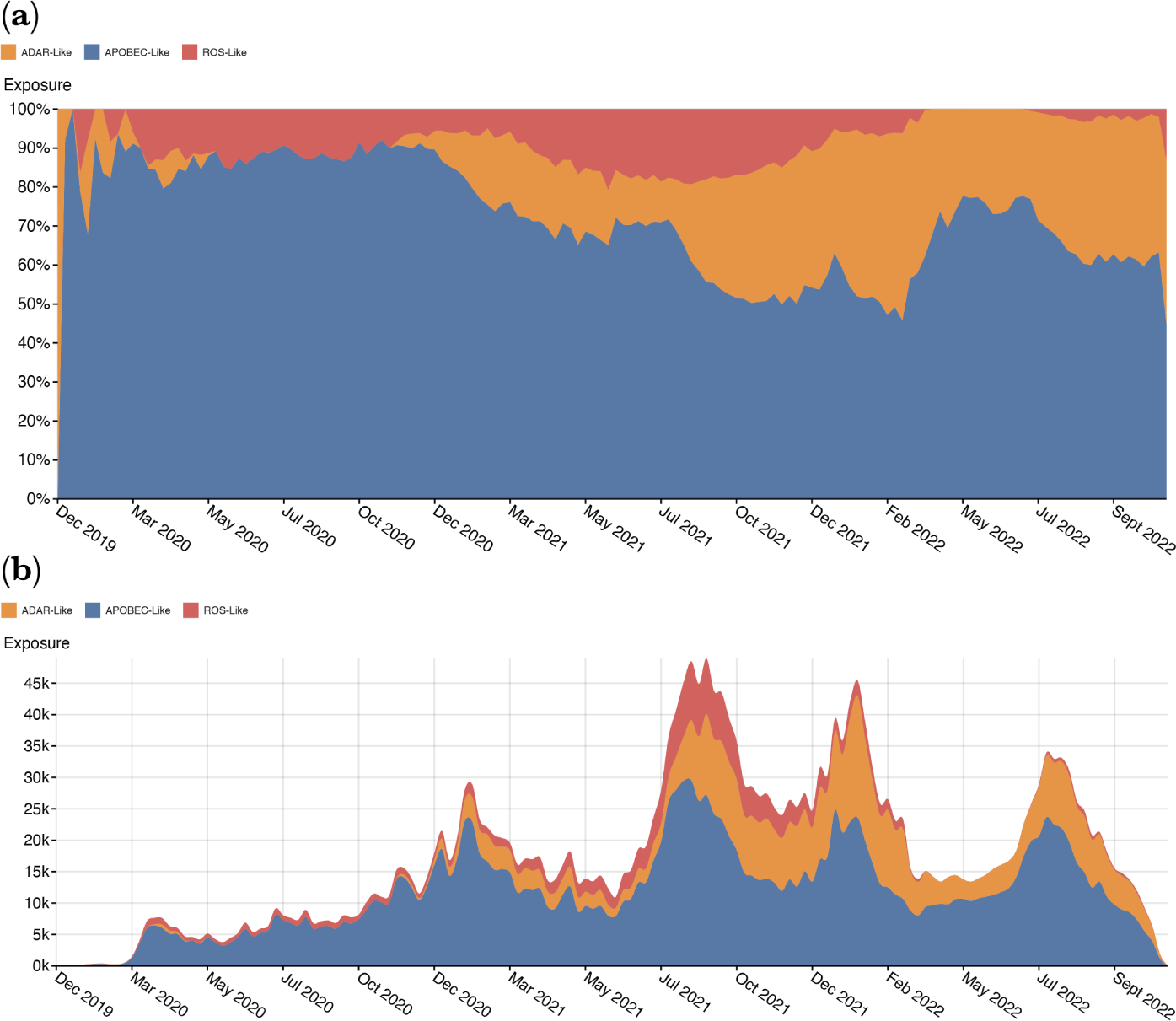
Signature Exposure plots showing the activities of the extracted mutation signatures over the duration of the pandemic. (a) shows the percentage activity of the signatures during a given week of the pandemic, with each colour representing a different signature. (b) shows the signature activities as their absolute values at each epidemic week.

**Fig. 5.**
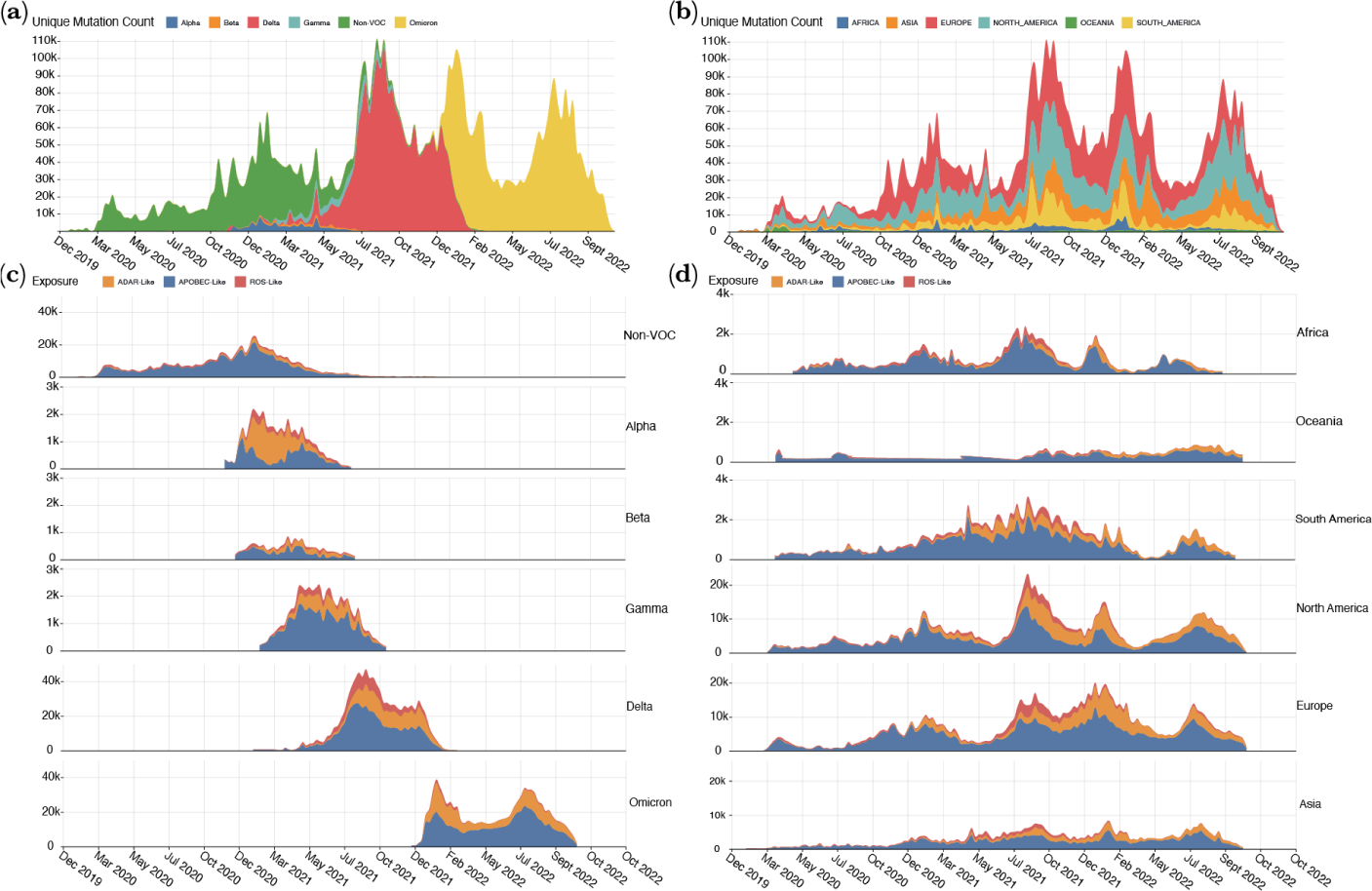
(a) Counts of unique mutations each epidemic week, with colours representing which continent the mutations came from. (b) Counts of unique mutations per week that are part of the mutational signature substitution-context features (i.e no gap mutations included). Each bar represents mutation counts at each epidemic week, with colours representing which lineage/group of lineages the mutations belong to. (c) Ridgeline plot showing the exposure of mutational signatures in SARS-CoV-2 lineage subsets. Exposures are coloured by the signature they have been attributed to. (d) Ridgeline plot showing the exposure of mutational signatures in SARS-CoV-2 continent subsets.

**Fig. 6.**
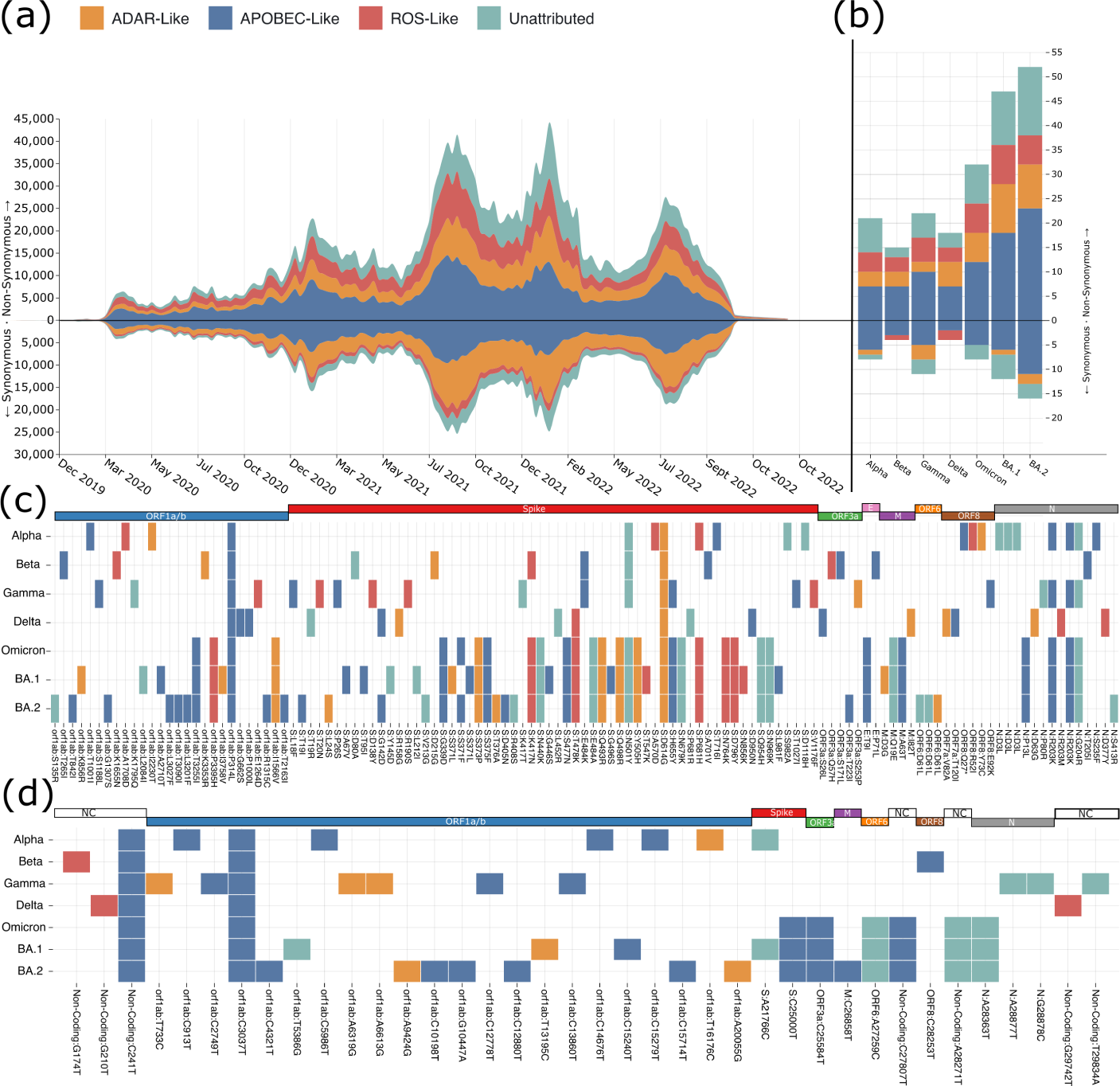
(a) Counts of unique mutations per week. Synonymous counts are on the bottom of the x-axis, while Non-synonymous counts are on the top. Each bar represents mutation counts at each epidemic week, with colours representing which process the mutations are attributed to. (b) Non-Synonymous and Synonymous mutations in the pseudo-references of identified variants of concern. APOBEC-like processes produce the majority of both synonymous and non-synonymous changes in all lineages. ROS-like changes are more often non-synonymous in the lineages of concern, with most lineages having no ROS-like synonymous changes. ADAR-like non-synonymous changes appear to have increased in the Omicron lineages, namely in BA.1 and BA.2. (c) Variant of concern amino acid changes coloured by the putative mutational process that caused the change. (d) Variant of concern synonymous nucleotide mutations coloured by the putative mutational process that caused the change.

**Fig. 7.**
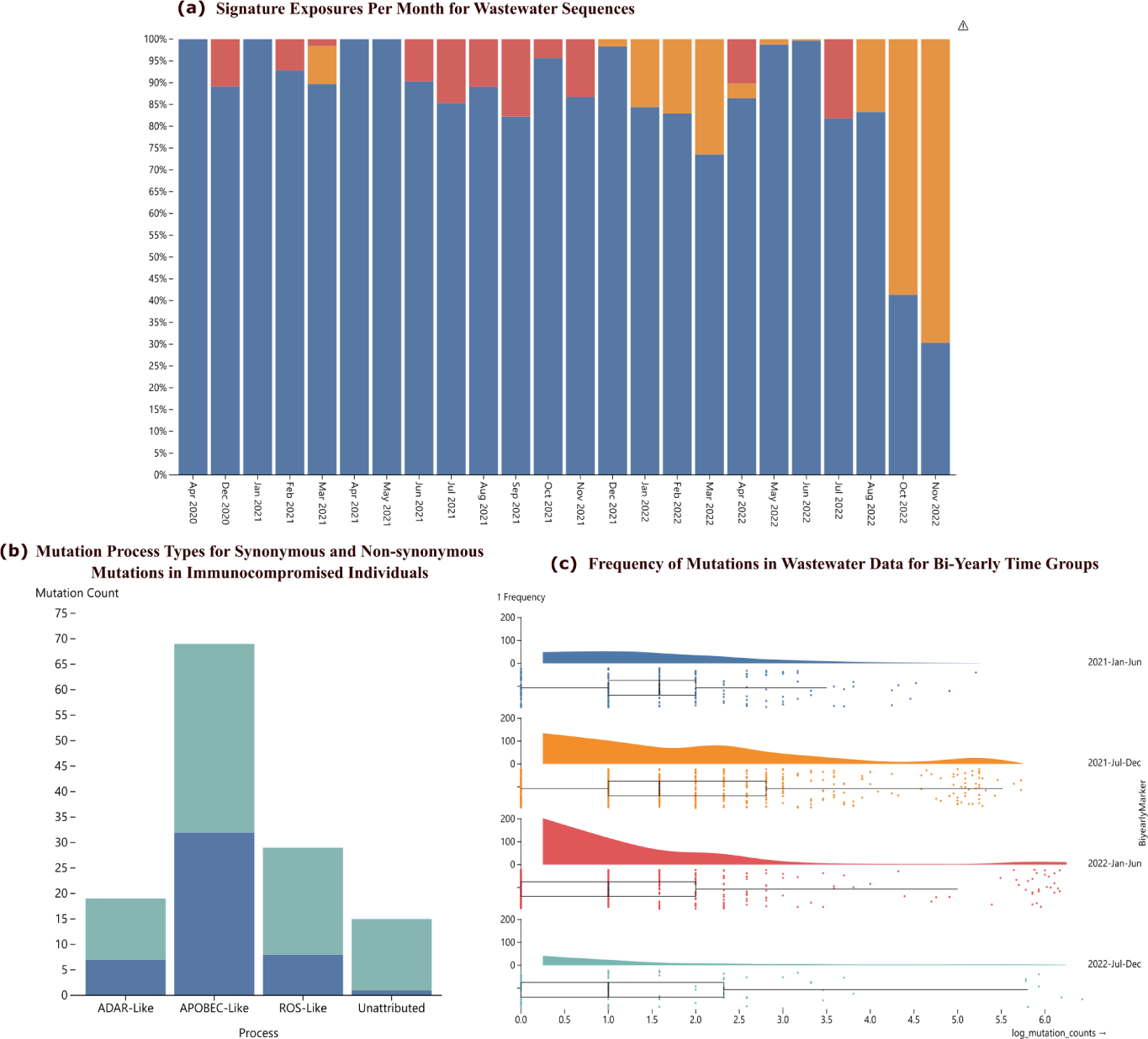
(a) Mutation counts in wastewater sequences for bi-yearly time groups. Highly mutated sequences cluster to the right especially during the 2021 July-December time group, as would be expected in this time of emerging omicron. (b) Signature exposures per month show similar trends in mutational processes as global data, although at a lower resolution and, interestingly, with a lower ADAR signature. (c) Mutations in consensus sequences from immunocompromised individuals contain mutation types corresponding to the global signatures. Mutation counts are presented as log10 of raw mutation counts. Interestingly, there are more synonymous mutations than in the global data, however the sample size is too small for this result to be conclusive.

The entry mechanism of the virus could impact the effect of ROS on the viral RNA since the two entry mechanisms take the viral RNA to the cytoplasm in different ways which could change if or when they might interact with ROS. The cell type preference of Omicron could also be involved since it is possible cells in the upper and lower respiratory tract have different quantities of ROS or different mechanisms for regulating it. Endosomal entry is favoured in the cells of the upper respiratory tract due to reduced expression of TMPRSS2 compared to lower respiratory tract cells. Omicron also shows lower replication efficiency in these lower respiratory cells, so fewer viral RNA’s are likely to be exported for forward transmission and/or sequencing. Delta on the other hand has a higher level of ROS-like exposure (Figure 5b) and has been found to both replicate better than Omicron in lung epithelial cells and have a 4x increased level of infection in cells with high expression of TMPRSS2 [43]. Delta is also known to fuse with the cell membrane considerably faster than prior VOCs [45], and like prior lineages of SARS-CoV-2 can form syncytia (cells fused together to form large multinucleated single cells). Omicrons excessive mutations in spike appear to have removed its ability to produce these syncitia which also may have implications in its mutation signature composition relative to previous variants. Any of these hypotheses would require further investigation via lab work to understand if Omicrons phenotypic differences result in the signature shift we observe, and whether it is in fact ROS causing these G→T changes. The G→T signature itself seems to display a context preference of TpG and ApG nucleotides, which could mean that the signature is in fact some other as yet unknown editing mechanism on the viral RNA rather than the ROS which is unlikely to have such a target preference. Signature ”switching” from APOBEC to ADAR changes happens from December 2020 onwards in the global dataset and appears consistently in the VOC and Continent subsets around this time point as well. Alpha experiences a major shift to ADAR-like mutations early in its time as the predominant VOC, although the APOBEC-like signature returns as the dominant set of changes towards the end of Alphas wave of infections. The Non-VOC subset appears to be the least impacted by ADAR-like changes, although this can mostly be explained by the number of Non-VOC sequences quickly declining after the emergence of the VOC lineages. Delta experiences a dramatic increase in ADAR-like and ROS-like exposure from July 2021, with ADAR-like exposure becoming the predominant signature towards the end of Deltas wave. The ADAR-like changes continue into Omicrons introduction, although it does decrease after the initial BA.1 wave from December 2021 to March 2022. It seems clear that while APOBEC-like mutations have dominated the mutational capacity of SARS-CoV-2 throughout the pandemic, this is beginning to change. It is possible that shifting activities are evidence of changing interactions between the viruses and the immune systems of the hosts they circulate within. Changes in population immunity via vaccination or previous infection may change the mutations that we observe in final reference sequences. Changing mutational process activity is unlikely to reflect the true activity of each process, but they are much more likely to show which processes are contributing mutations that eventually make it into circulating viruses. This is something that would be missed by intra-patient samples, although these are much more likely to give a better idea of true mutational process activity.

All lineages of concern we assessed show predominantly non-synonymous mutations, and all putative mutational signatures produced more non-synonymous changes than synonymous changes. Synonymous changes were much more likely to occur in ORF1a/b, which would be expected due to its size as the largest ORF, but this pattern is not observed with non-synonymous mutations which are mainly centred on the spike protein (6c,d). This is consistent with spike being under intense immune pressure since it is the main glycoprotein for SARS-CoV-2. Spike elicits much of the antiviral response which is why it is the protein used in the SARS-CoV-2 vaccines. As such, spike is under greater pressure to change in order to escape the host immune response while maintaining its function of binding to host cells. APOBEC-like changes are the predominant source of mutations in all SARS-CoV-2 VOCs that we analysed, followed bu unattributed mutations, ADAR-like changes and the ROS-like changes. ROS-like changes were unlikely to be synonymous with only Beta and Delta containing such changes (6d). This is also reflected in the global unique mutation count where ROS-like mutations only represent a small fraction of synomymous changes. ADAR-like mutations appear the most likely to be synonymous (6a) but this does not seem to be observed in the VOC lineages where most ADAR-like changes are non-synomymous (6b) except for in the Gamma VOC.

Each putative mutational process has produced important changes within SARS-CoV-2. Given the virus now circulates freely in the population with limited restrictions it seems likely that this will continue. Mutational signatures offer a way of tracking these changes over time and may provide insights into the interplay between intrinsic viral properties and mutational processes effects on the virus as it continues to accumulate more change.

## 4 Methods

### 4.1 Data

The findings of this study are based on metadata associated with 13,281,213 sequences available on GISAID up to October 26, 2022, and accessible at doi.org/10.55876/gis8.221201qs. Sequences were filtered to remove records of non-human hosts, lengths less than 20,000 nucleotides, non-assigned lineages, greater than 30% unknown bases, sequences collected before 24/12/2019 and those with excessive mutations/deletions.

Publicly available daily SARS-CoV-2 cases, tests performed and total vaccinations per capita were obtained from OWID [46]. Country-level government stringency indices were downloaded from OxCGRT [47]. Government stringency indices are composed of nine indicators: school closure, workplace closure, cancellation of public events, stay at home order, public information campaigns, restrictions on public gatherings, public transport, internal movement and international travel. The index on a given day ranges 0 to 100 and is calculated as the mean of the nine indicators, with higher indices indicating stricter regulations. If responses vary at sub-national levels, the index at the strictest level is used [47].

Wastewater findings of this study are based on metadata associated with 1,343 sequences available on GISAID and accessible at doi.org/10.55876/gis8.230406qg. Wastewater sequences were downloaded from the ‘wastewater data’ section of GISAID in December 2022.

Sequences for immunocompromised individuals were downloaded from GISAID in November 2022. Analysis of this was based on the meta-data associated with 34 sequences available on GISAID and accessible at doi.org/10.55876/gis8.230406fb. Sequences werechosen based on the known list of sequences used in [35]. Sequences were aligned to the COVID reference genome before use.

### 4.2 Design

Predictors of SARS-CoV-2 reported cases were explored using a linear model at both country and continent levels. We collected continuous dependent variables that changed with every calendar date did not remain constant for finite amounts of time. These were classified into two groups: (i) public health measures (government stringency, testing capacity and vaccination) and (ii) viral properties (diversity and fitness). We examined the data for completeness of predictive variables. Missing vaccination data was handled as no (zero) vaccinations were administered. With exception of vaccinations, variables with less than 70% of the countries reporting data were not included. The number of SARS-CoV-2 diagnostic tests performed was excluded as a predictor due to missing data. We defined virus fitness as the sum of previously identified [48] amino acid substitutions that increase SARS-CoV-2 fitness divided by the sum of total genomes and the log of total mutations [48].

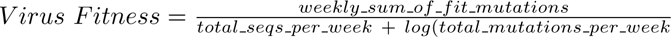

Diversity was calculated by dividing distinct lineages by the total number of genomes in a given week. Sequences reported in GISAID were assumed to be representative of the diversity of infections for that continent/country.

### 4.3 Linear Model

For continent-level analysis, data from all contributing countries was used to fit the linear model. Data was smoothed using a 14-days rolling average to limit possible noise and identify simplified changes over time. Continental government stringency index was calculated as the daily average of country-level indices. Pearson’s correlation was used to test for correlation among the variables. Multiple linear regression was fitted to evaluate the relationship between infection rate (daily cases per capita) as the outcome and the public health measures and viral properties as predictors within the different continents. The regression models were fitted on data from 01 April 2020 onwards, as (sequence) data addition remained constant after this. The country-level analysis was carried out for countries with less than 50 days of missing genome data using a similar approach.

### 4.4 Pandemic Plots

Case numbers and sequence data were aggregated by their respective continents, a 14-day rolling average was used to smooth out daily infection rates and categorical variables were summarised by counts. Proportions of lineages were calculated in 14-days bins and the most common lineages were visualised per continent.

### 4.5 Tree-based Referencing

The rapid spread and evolution of SARS-CoV-2 means that most viral sequences currently circulating are mutationally distinct from the early pandemic reference genome Wuhan-Hu-1 [49]. Continuing to count mutations against the early reference sequence can result in mutations being allocated the wrong substitution category (i.e., A→T instead of a C→T) where sites have mutated multiple times.

[37] tackled this issue by building a tree of clustered sequences to remove ancestral mutations, however, we can utilise the available SARS-CoV-2 tree generated as part of the Pangolin [8] nomenclature to generate ancestral sequences. This means that sequences from the lineage B.1 are compared against a generated reference sequence for the B lineage rather than the Wuhan-Hu-1 sequence.

Reference sequences were generated for each of the Pangolin lineages in the alignment. A nucleotide was included in the Pangolin reference if it exceeded a frequency threshold of greater than 75% of the samples from the lineage. If this threshold was not reached, the reference nucleotide of the nearest parental lineage was used. Building intermediate references also meant that counting inherited mutations could be avoided. Since mutations were identified relative to their nearest parental Pangolin lineage, mutations inherited are not counted since relative to this sequence there hasn’t been a mutation.

### 4.6 Pseudo-Sampling

Mutations were binned into categories composed of their substitution type (e.g cytosine → thymine = CT) and their mutation context. The mutation context is the mutated base and the nucleotides at the 5’ and 3’ positions of the mutated base. There are a total of 192 types of substitution-context matchings that can appear (12 possible single nucleotide changes x 4 possible nucleotide 5’ x 4 possible nucleotide 3’). Every sequence produces a single count vector of mutation category counts, with the total count matrix becoming the mutational catalogue of the virus. SARS-CoV-2 genome sequences from any one week of the epidemic often have very few newly-arisen unique mutations. Extracting mutational signatures for such low mutation counts is not advisable and is unlikely to produce meaningful results. To resolve this problem, we define each sample as a time-point (Epidemic Week) and decompose signatures from the counts at each time-point rather than from each sequence. This shrinks the mutational catalogue of the virus from millions of samples down to less than 200 samples for each Epidemic Week.

### 4.7 Non-Negative Matrix Factorisation

NMF(Non-negative matrix factorisation)[50] was used to split the mutational catalogue into 2 sub matrices. One matrix represents the mutational signatures, the other matrix represents the exposure of the signatures. These matrices were used to reconstruct the original mutational catalogue with some degree of error. To verify the validity of the identified signatures, NMF was performed 100 times for each value of N, with N representing the number of signatures to extract from the mutational catalogue. For each iteration, a new mutational catalogue was generated using bootstrap re-sampling of the original matrix and removal of any mutational categories that did not account for more than 0.5% of mutations. The signatures were then clustered together using K-Means, with the cluster means forming the new signatures. Clusters were then assessed using the Silhouette Score to determine the quality of the clustering. Clusters with high silhouette scores are well separated from other clusters and are dense and well formed. Cosine similarity was used to determine if the signature was reliably extracted from the cluster. The cosine similarity was calculated between signatures extracted from the whole mutational catalogue and the cluster means of the signature clusters. A higher cosine similarity indicates that the cluster mean shows a similar pattern to the initial mutational signature. An N value of 3 was selected due to the reduction of the reconstruction error plateauing around 3, and the marked decrease in silhouette score for signatures greater than 3. The average cosine similarity between signatures and clusters was consistently above 0.95 for each cluster and had an average of 0.98 for all 3 clusters when clustering was repeated 100 times. Signatures can therefore be reliably extracted from the bootstrapped catalogues, are robust and thus are unlikely to be artefacts.

### 4.8 Non-Negative Least Squares Regression

A Non-Negative Least Squares (NNLS) Regression was used to produce positive exposure weights for each of the signatures in each of the datasets. The non-negativity of the regression ensures that the weights like the signatures continue to represent an additive process. The NNLS weights can then represent the exposures of the signatures on each dataset.

### 4.9 Consensus Lineage and Continent Signatures

Mutational catalogues were constructed for each continent and each of the Variant of Concern (VOC) lineages (Alpha, Beta, Gamma,Delta, Omicron). The global signatures were then used to extract exposures for each of the mutational catalogues to determine how processes varied between each mutational catalogue subset. VOC sequence sets were filtered so that weeks with fewer than 100 sequences were excluded.

## Acknowledgments

We gratefully acknowledge all data contributors, i.e., the authors and their originating laboratories responsible for obtaining the specimens, and their Submitting laboratories for generating the genetic sequence and metadata and sharing via the GISAID Initiative, on which this research is based. The authors acknowledge funding from the Medical Research Council, (MRC, MC UU 12014/12 and a Doctoral Training Programme in Precision Medicine studentship to KDL), the UK Department for International Development (DFID) under the MRC/DFID Concordat agreement (MC PC 20010), the Biotechnology and Biological Sciences Research Council (BBSRC, BB/V016067/1), Engineering and Physical Sciences Research Council (EPSRC, EP/R018634/1), European Union’s Horizon 2020 research and innovation programme project PANCAIM (101016851) and the Wellcome Trust (220977/Z/20/Z).

We would also like to thank Spyros Lytras, Francesca Young, Sejal Modha, Andres Gomez and Procheta Sen for their helpful comments throughout the process of writing and preparing this manuscript.

## Appendix A

**Fig. A1.**
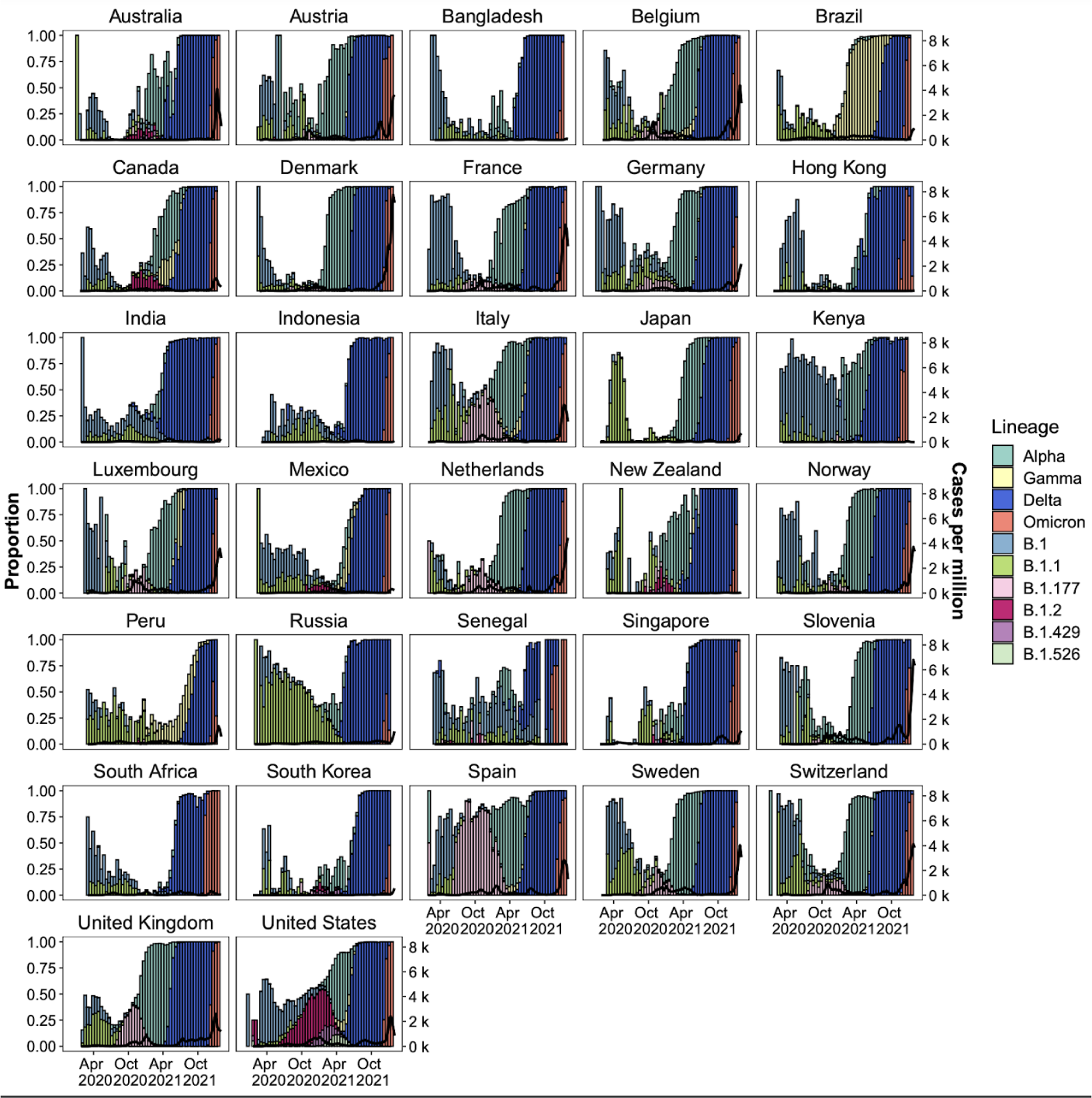
Country-level SARS-CoV-2 lineage dynamics. Solid bars show the biweekly proportions of the top five lineages per country. Bars are coloured by lineage. Countries included in this analysis based on temporal data completeness.

**Fig. A2.**
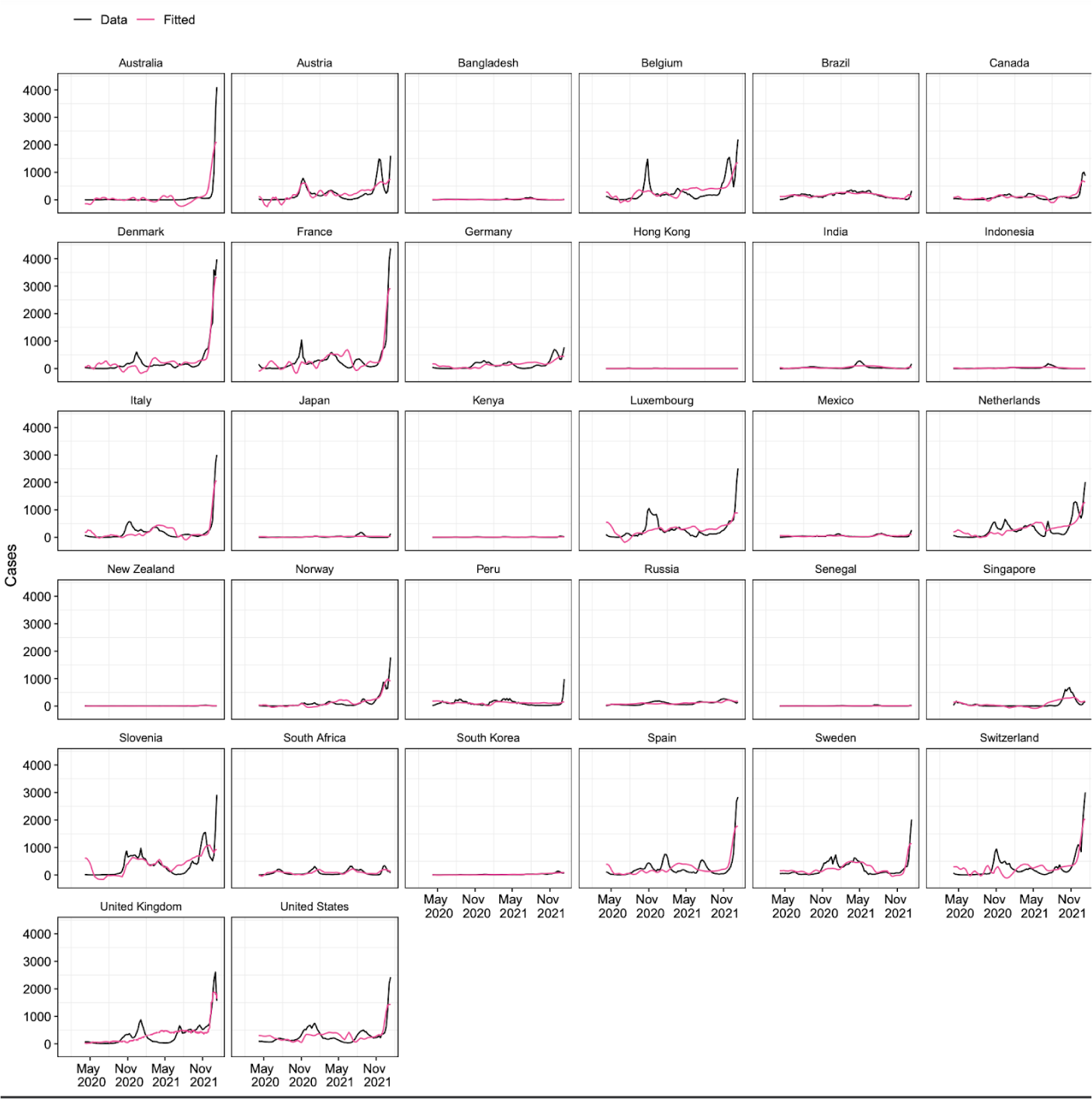
Model-fitting of country-level SARS-CoV-2 reported cases. Black solid lines show a 14-day rolling average of reported SARS-CoV-2 cases. Pink solid lines show fitted mean response values of infection rates with predictor values as input. Countries included in this analysis based on temporal data completeness.

**Fig. A3.**
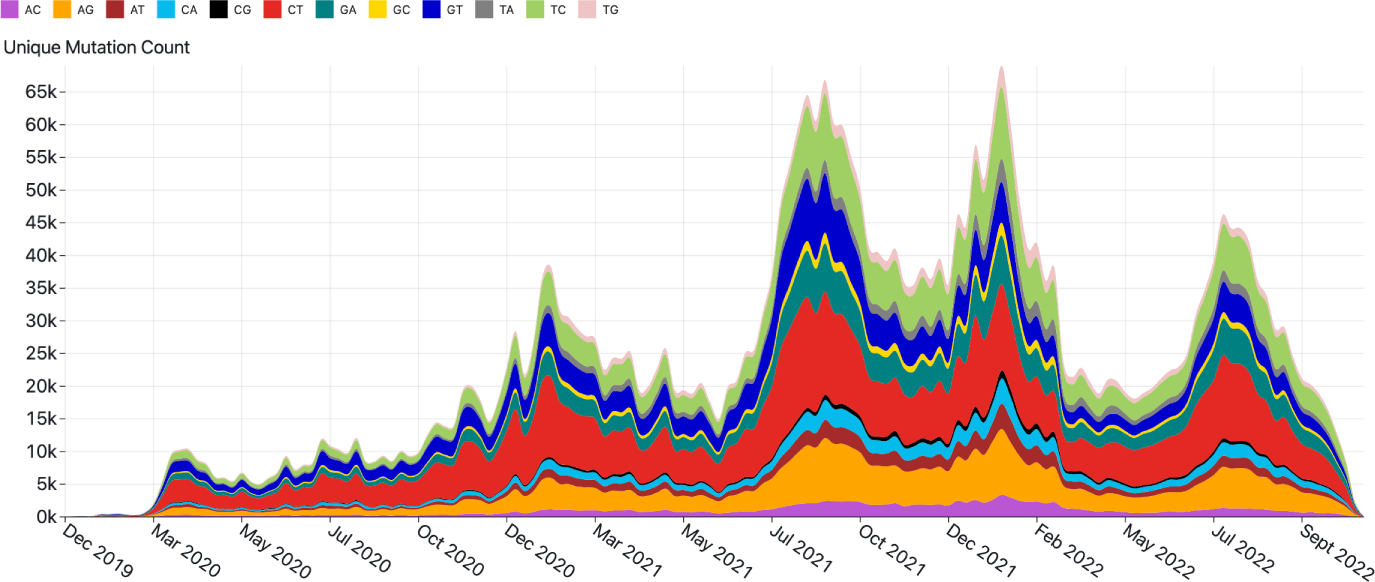
Counts of unique substitutions per week of the pandemic. Bars are coloured by substitution category.

**Table A1.** Effect of public health measures (government stringency and vaccination) and viral properties (diversity and fitness) on infection rates.

|  |  | Africa | Asia | Europe | North America | South America | Oceania |
| --- | --- | --- | --- | --- | --- | --- | --- |
| Intercept | Estimate | 19.36 | 39.07 | 14.03 | 813.08 | 10.91 | 431.03 |
|  | S.E | 2.63 | 7.95 | 54.78 | 39.83 | 1.15 | 32.62 |
|  | t value | 7.36 | 4.92 | 0.26 | 20.41 | 9.47 | 13.21 |
|  | Pr(>t) | 7.51E-13 | 1.18E-06 | 0.8 | <2.00E-16 | <2e-16 | <2.00E-16 |
| Govt stringency | Estimate | -0.24 | -0.47 | 1.17 | -10.92 | 0 | -3.47 |
|  | S.E | 0.03 | 0.11 | 0.57 | 0.67 | 0.01 | 0.39 |
|  | t value | -7.07 | -4.25 | 2.03 | -16.24 | 0.14 | -8.99 |
|  | Pr(>t) | <4.99E-12 | 2.57E-05 | 0.04 | <2.00E-16 | 0.89 | <2.00E-16 |
| Vaccination | Estimate | 2.43 | -0.12 | -0.96 | 1.66 | 0.61 | -1.97 |
|  | S.E | 0.2 | 0.06 | 0.32 | 0.36 | 0.02 | 0.16 |
|  | t value | 12.25 | -2.08 | -2.98 | 4.56 | 38.23 | -12.12 |
|  | Pr(>t) | <2.00E-16 | 0.04 | 3.01E-03 | 6.38E-06 | <2e-16 | <2.00E-16 |
| Virus fitness | Estimate | 1.84 | 50.44 | 145.76 | -343.32 | -9.26 | 159.13 |
|  | S.E | 1.24 | 4.15 | 23.97 | 28.61 | 0.97 | 14.21 |
|  | t value | 1.49 | 12.15 | 6.08 | -12 | -9.56 | 11.19 |
|  | Pr(>t) | 1.40E-01 | <2.00E-16 | 2.34E-09 | <2.00E-16 | <2e-16 | <2.00E-16 |
| Virus diversity | Estimate | 1.35E+00 | 15.01 | 125.91 | -291.74 | -29.56 | -181.72 |
|  | S.E | 4.53E+00 | 17.52 | 139.74 | 80.24 | 2.92 | 54.59 |
|  | t value | 3.00E-01 | 0.86 | 0.9 | -3.64 | -10.11 | -3.33 |
|  | Pr(>t) | 7.70E-01 | 0.39 | 3.70E-01 | 3.05E-04 | <2e-16 | 0.00034 |
| Residual S.E |  | 218.9 on 147 DF | 15.85 on 512 DF | 95.91 on 512 DF | 86.19 on 512 DF | 851.8 on 512 DF | 47.84 on 512 DF |
| Multiple R squared |  | 0.63 | 0.56 | 0.17 | 0.42 | 0.87 | 0.66 |

Abbreviations: S.E = Standard error DF = degrees of freedom Govt = Government Data: Our World in Data [26], Oxford Covid-19 Government Response Tracker [27] and GISAID(www.gisaid.org).

**Table A2.** Proportion of common lineages/variants globally

| Lineage | Sum | Proportion |
| --- | --- | --- |
| Delta | 4,087,909 | 0.563 |
| Alpha | 1,150,798 | 0.158 |
| Omicron | 647,553 | 0.089 |
| B.1.2 | 127,557 | 0.018 |
| Gamma | 121,393 | 0.017 |
| B.1 | 111,849 | 0.015 |
| B.1.177 | 74,643 | 0.010 |
| Beta | 40,786 | 0.006 |
| B.1.1.214 | 18,160 | 0.002 |
| D.2 | 13,340 | 0.002 |
| B.1.621 | 11,050 | 0.002 |
| B.1.1.284 | 9,334 | 0.001 |
| C.37 | 9,287 | 0.001 |
| Total | 6,423,659 | 0.884 |

**Table A3.** Correlation between infection rate and predictor variables across different continents

| Continent | Infection rate | Virus fitness | Virus diversity | Stringency index | Vaccination |
| --- | --- | --- | --- | --- | --- |
| Africa | 1 | 0.55 | -0.65 | -0.59 | 0.73 |
| Asia | 1 | 0.73 | -0.65 | -0.23 | 0.5 |
| Europe | 1 | 0.28 | -0.19 | 0.2 | 0.01 |
| North America | 1 | -0.15 | -0.01 | -0.39 | -0.05 |
| Oceania | 1 | 0.59 | -0.68 | -0.2 | 0.91 |
| South America | 1 | 0.72 | -0.6 | -0.57 | 0.23 |

